# Inhibition of the phosphoinositide 3-kinase-AKT-cyclic GMP-c-Jun N-terminal kinase signaling pathway attenuates the development of morphine tolerance in a mouse model of neuropathic pain

**DOI:** 10.1101/2020.10.14.340067

**Authors:** T. Okerman, T. Jurgenson, M. Moore, A. H. Klein

## Abstract

**Background:** Opioid management of chronic pain can cause opioid-induced analgesic tolerance and hyperalgesia, complicating clinical pain-management treatments. Research presented here sought to determine if opioid induced tolerance is linked to activity changes within the PI3Kγ-AKT-cGMP-JNK intracellular signaling pathway in spinal cord or peripheral nervous systems.

**Methods:** Morphine or saline injections were given subcutaneously twice a day for five days (15 mg/kg) to male C57Bl6 mice. A separate cohort of mice received spinal nerve ligation (SNL) one week prior to the start of morphine tolerance. Afterwards, spinal cord, dorsal root ganglia, and sciatic nerves were isolated for quantifying total and phosphorylated-JNK levels, cGMP, and gene expression analysis.

**Results:** Gene expression for the PI3Kγ-AKT-cGMP-JNK signaling pathway including, *Akt1, Akt2, Akt3, Pik3cg*, *Pten, Jnk3*, and *nNos1* were decreased in the spinal cord with varied expression changes in the dorsal root ganglia and sciatic nerve of morphine tolerant and morphine tolerant mice after SNL. We observed significant increases in total and phosphorylated-JNK levels in the spinal cord, total JNK in dorsal root ganglia, and cGMP in the sciatic nerve of morphine tolerant mice with SNL. Pharmacological inhibition of PI3K, nNOS, or JNK, using thalidomide, quercetin, or SP600125, attenuated the development of morphine tolerance in mice with SNL as measured by thermal paw withdrawal.

**Conclusions:** Overall, the PI3K/AKT intracellular signaling pathway is a potential target for reducing the development of morphine tolerance. Continued research into this pathway will contribute to the development of new analgesic drug therapies.

## Introduction

Neuropathic pain affects millions of patients (Gordon Smith and Robinson Singleton, 2006; Scholz et al., 2019), and while the efficacy of opioid medications for neuropathic pain is debatable, opioids are still used in up to 30% of patients experiencing neuropathic pain (DiBonaventura et al., 2017). When mu-opioid receptors (MOPs) are continuously stimulated through repeated opioid use, tolerance develops. Previous research suggests overexposure of the MOP to agonists initiates phosphorylation of the receptor via kinase activity, resulting in the recruitment of β-arrestin and downregulation of MOPs (Benovic et al., 1989). However, recent studies show inhibition of β-arrestin phosphorylation (Kliewer et al., 2019) or genetic deletion of β-arrestin 2 (Bohn et al., 2002) does not completely prevent the development of opioid tolerance and can potentially worsen opioid side effects.

ATP-sensitive potassium (K_ATP_) channels are inwardly-rectifying potassium channels found in the brain, dorsal root ganglia, and superficial dorsal horn of the spinal cord (Fotinou et al., 2013; Gerzanich et al., 2019; Zoga et al., 2010). K_ATP_ channels are downstream targets of opioid receptors (Cunha et al., 2010) and previous studies have reported a decrease in K_ATP_ expression in dorsal root ganglia after a painful nerve injury (Sarantopoulos et al., 2003) with or without underlying tolerance to morphine (Fisher et al., 2019). K_ATP_ channel agonists given concurrently with morphine lead to the attenuation of morphine tolerance in mice (Cao et al., 2016; Fisher et al., 2019). These findings suggest the activity of K_ATP_ channels in primary afferent neurons decrease after nerve injury and/or morphine tolerance and contribute to hypersensitivity. K_ATP_ channels are reportedly activated by at least two separate signaling pathways including the PI3K-AKT-nNOS-cGMP-PKG pathway (Cunha et al., 2010), and the ROS/Calmodulin/CaMKII signaling pathway (Chai et al., 2011). The PI3K-AKT-nNOS-cGMP intracellular signaling pathway contains several second messengers implicated in pain sensitization and are a promising target for studying mechanisms underlying opioid dependence.

Of the three known PI3K isoforms, the PI3Kγ variant is known to play a role in opioid receptor signaling and is highly expressed in the nervous system (Cunha et al., 2010). Previous reports show Akt inhibition within the PI3K/Akt/mTOR signaling pathway alleviates chronic neuropathic pain, as phosphorylated-Akt expression is increased in the spinal cord in rodent neuropathic pain models (Guedes et al., 2008; Guo et al., 2017). Activation of Akt and neuronal nitric oxide synthase (nNOS) in the periaqueductal grey is responsible for sustained potentiation of NMDAR and results in MOP inhibition and produces an analgesic tolerance (Sánchez-Blázquez et al., 2010). Conversely in the peripheral nervous system, morphine tolerance is attenuated by blocking the NOS-cGMP-PKG-JNK pathway in sciatic nerve injured mice (Hervera et al., 2012) and stimulation of downstream JNK signaling is thought to precipitate tolerance in some opioid tolerance models (Melief et al., 2010). The research presented here sought to determine if opioid induced tolerance is linked to increased or decreased activity in the PI3Kγ-AKT-cGMP-JNK intracellular signaling pathway in the peripheral versus central nervous system.

## Experimental Procedures

### Animals

Experimental procedures involving animals were performed in accordance with the US National Research Council’s Guide for the Care and Use of Laboratory Animals, the US Public Health Service’s Policy on Humane Care and Use of Laboratory Animals, and Guide for the Care and Use of Laboratory Animals. Protocols involving animals were approved by the University of Minnesota Institutional Animal Care and Use Committee. Adult C57Bl/6 WT male mice were acquired from Charles River Laboratories (Raleigh, NC) at 5-6 weeks old (21 ± 4.0 g). Mice were randomly split into three different experimental groups for protein, nucleotide, and gene expression analysis. Two groups of mice were administered saline or morphine (MT, 15 mg/kg) twice a day for five days (7-8 weeks old), and a third group of mice with spinal nerve ligation (SNL) one week before chronic morphine treatment (MT + SNL). A separate cohort of MT + SNL mice were split into two additional experimental groups for separate behavior studies treated with either vehicle or with a PI3K/AKT pathway inhibitor (below). At the end of the study, mice were euthanized with 5% isoflurane in oxygen and decapitated. Spinal cords (SC), dorsal root ganglia (DRG), and sciatic nerves (SN) were isolated, snap frozen in liquid nitrogen and immediately stored at -80°C.

### Drugs and Delivery

Vehicle (100 µL saline, subcutaneous) treated mice were used as controls for morphine tolerant mice without SNL and morphine tolerant mice after SNL. Morphine (Sigma Chemical, St. Louis, MO) in saline was administered twice per day for five days (15 mg/kg, s.c.). For MT+SNL mice treated with or without PI3K/AKT pathway inhibitors, vehicle (20% DMSO, 5% Tween 20, in saline, 100 µL, i.p.) controls or drug-treated animals received injections twice per day for five days. Quercetin (SC-206089A, 60 mg/kg, 100 µL, Santa Cruz Biotechnology, Dallas TX) in saline was administered 30 minutes prior to morphine. Thalidomide (J60271, 100 m/kg, 100uL, Alfa Aesar, Ward Hill, MA) in 10% DMSO in saline was administered 15 minutes prior to morphine. SP600125 (S-7979, 10 mg/kg, 100uL, LC Laboratories, Woburn, MA) in 20% DMSO and 5% Tween 20 in saline was administered 30 minutes prior to morphine. Timing of inhibitor injections relative to morphine was based off previous studies (Filho et al., 2008; Hassanzadeh et al., 2016; Marcus et al., 2015).

### Spinal Nerve Ligation

To model the effects of neuropathic pain in a mouse model, spinal nerve ligation was performed (Masuda et al., 2017). Mice were anesthetized with 3% isoflurane (1.5% post-anesthetization) in an O_2_ carrier. The area above the L3 to S3 vertebrae was shaved and cleaned with povidone iodine (Betadine,1413, Dynarex, Orangeburg, NY). An incision into the skin and fascia was made above the L5 to S1 vertebra. To expose and transect the left L4 spinal nerve, the paravertebral musculature was removed from the vertebral transverse processes, along with a portion of the L5 transverse process. The incision was closed with two layers of 4-0 vicryl sutures (J304H, Ethicon, Bridgewater, NJ). The right spinal nerves were left intact (contralateral). Fourteen days elapsed before behavioral testing without perioperative analgesics.

### Thermal Paw Withdrawal Latency

Mice were acclimated to the testing apparatus at least 3 days prior to formal testing at 7-8 weeks old for saline and MT mice and 10 weeks old for MT + SNL mice. The testing environment consisted of individual acrylic containers on a glass pane heated to 30°C. The Modified Hargreaves’ method was used to measure thermal paw withdrawal latency (TPW, 390G, IITC, Woodland Hills, CA) (Banik and Kabadi, 2013). TPW was determined by the amount of time, is seconds, required to elicit a nocifensive response after a radiant heat beam was placed under the plantar surface of the hindpaw. Baseline measurements were collected five times for each hind paw before any drug administration and a single TPW measurement at 0, 30, and 60 minutes post morphine administration. Measurements were taken before the start of morphine tolerance and on days 1, 3, and 5.

### mRNA Isolation and Quantitative PCR

Total RNA (50 ng) was isolated from vehicle, MT, and MT+SNL mice from tissues collected on day 6 (post behavioral testing) using Tri Reagent (T9424, Sigma Aldrich, St. Louis, MO) and RNeasy Micro kit (74106, Qiagen, Valencia, CA) with DNase digestion steps according to manufacturer’s protocol. A cDNA library was created form each mRNA sample using the Omniscript RT kit (205113, Qiagen, Valencia CA) with random nonamers (Integrated DNA Technologies, Coralville, Iowa). Quantitative PCR was performed using a SYBR Green master mix (Roche Diagnostics, Indianapolis, IN) on a Roche LightCycler 480 System (Roche Diagnostics, Indianapolis, IN) with. For every gene, the cDNA copy number was quantified against a 6 point, 10-fold serial dilution (5e^7^ to 5e^2^) cDNA standard. Expression of the housekeeping gene *Rn18s* was compared across samples to check for comparative cDNA amplification of each sample. RT negative samples were ran as an internal control. Table 1 provides a list of primers used in this study.

**Table 1.**
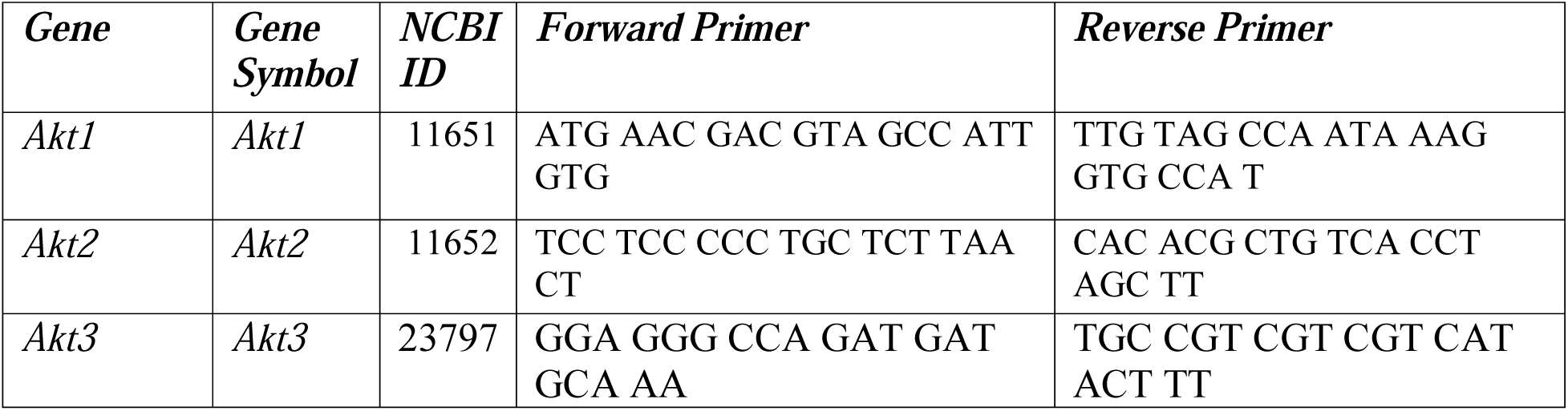
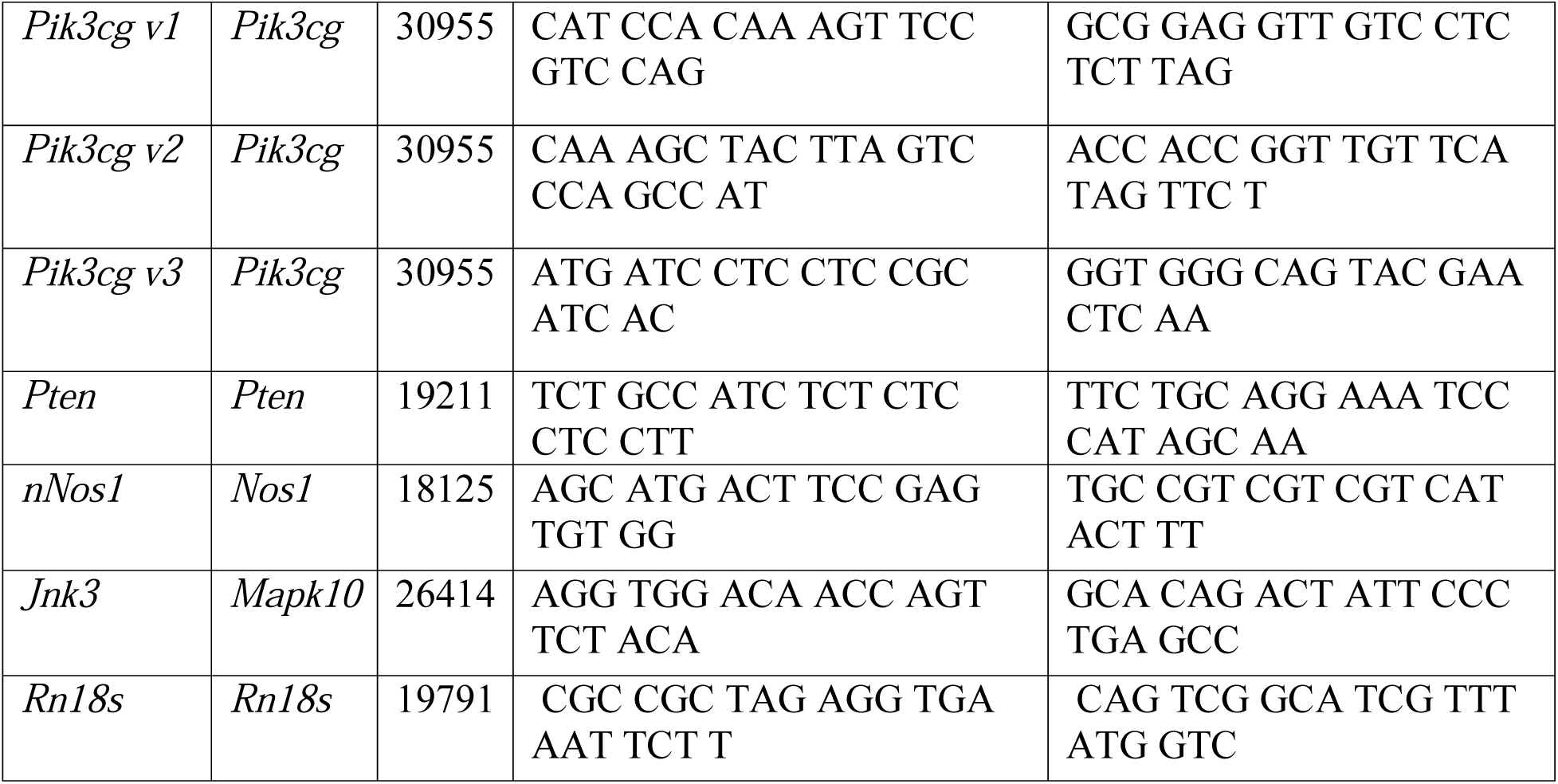
List of forward and reverse qPCR primers. “v” indicates splice variants.

### ELISA

cGMP, pJNK and total JNK levels were compared across vehicle, MT, and MT+SNL treatment groups and were measured using ELISA kits according to manufacturer’s protocol. cGMP assays (581021, Cayman Chemical Company, Ann-Arbor, MI) were scaled to accommodate tissues <0.05 g and were allowed two hours to develop on an orbital shaker before reading at 410 nm. pJNK and total JNK (ab176662, Abcam, Cambridge, MA) assays were read at 450 nm. Assays were read using a BioTek Synergy 2 Plate Reader (BioTek Instruments, Inc., Winooski, VT).

### Data Analysis

The appropriate one-way, two-way, or repeated measures ANOVA was used to determine the significance of treatment groups for gene expression, TPW, and ELISAs followed by a Dunnett post-hoc test with saline treatment as the control. The data are presented as means ±SEM, and p< 0.05 considered statistically significant. A list of ANOVA statistics are listed in Table 2. All statistics were completed using Prism 6 (GraphPad Software, La Jolla, CA).

**Table 2.** ANOVA statistics for main effects and interactions.

| Figure | Source | Fold $\Delta$ | df | F | p | CI | Post-hoc |
| --- | --- | --- | --- | --- | --- | --- | --- |
| 2A | *Pik3cg v1* Saline vs Ipsi | 0.043 | 3, 16 | 169.5 | <0.0001 | 596.8 to 847.3 | Dunnett |
|  | *Pik3cg v1* Saline vs Contra | 0.033 | 3, 16 | 169.5 | <0.0001 | 604.5 to 855.0 | Dunnett |
|  | *Pik3cg v2* Saline vs Ipsi | 0.005 | 3, 16 | 17.97 | <0.0001 | 12514 to 44096 | Dunnett |
|  | *Pik3cg v2* Saline vs Contra | 0.002 | 3, 16 | 17.97 | <0.0001 | 12607 to 44189 | Dunnett |
|  | *Pik3cg v3* Saline vs Ipsi | 0.027 | 3, 16 | 14.43 | <0.0001 | 1346 to 5494 | Dunnett |
|  | *Pik3cg v3* Saline vs Contra | 0.013 | 3, 16 | 14.43 | <0.0001 | 1398 to 5546 | Dunnett |
|  | *Akt1* Saline vs Ipsi | 0.071 | 3, 16 | 13.47 | 0.0001 | 13463 to 40015 | Dunnett |
|  | *Akt1* Saline vs Contra | 0.057 | 3, 16 | 13.47 | 0.0001 | 13865 to 40418 | Dunnett |
|  | *Akt2* Saline vs Ipsi | 0.019 | 3, 16 | 25.76 | <0.0001 | 2137 to 6459 | Dunnett |
|  | *Akt2* Saline vs Contra | 0.014 | 3, 16 | 25.76 | <0.0001 | 2160 to 6482 | Dunnett |
|  | *Akt3* Saline vs Ipsi | 0.013 | 3, 16 | 29.33 | <0.0001 | 22649 to 54184 | Dunnett |
|  | *Akt3* Saline vs Contra | 0.009 | 3, 16 | 29.33 | <0.0001 | 22821 to 54356 | Dunnett |
|  | *nNos1* Saline vs Ipsi | 0.020 | 3, 16 | 33.41 | <0.0001 | 12842 to 29331 | Dunnett |
|  | *nNos1* Saline vs Contra | 0.020 | 3, 16 | 33.41 | <0.0001 | 12856 to 29344 | Dunnett |
|  | *Jnk3* Saline vs Ipsi | 0.033 | 3, 16 | 20.15 | <0.0001 | 16620 to 67781 | Dunnett |
|  | *Jnk3* Saline vs Contra | 0.020 | 3, 16 | 20.15 | <0.0001 | 17193 to 68353 | Dunnett |
| 2B | *Pik3cg v2* Saline vs Contra | 0.367 | 3, 16 | 4.948 | 0.0128 | 722.1 to 34772 | Dunnett |
|  | *Pik3cg v3* Saline vs Morphine | 1.525 | 3, 16 | 98.53 | <0.0001 | -4805 to -1558 | Dunnett |
|  | *Pik3cg v3* Saline vs Ipsi | 0.066 | 3, 16 | 98.53 | <0.0001 | 4037 to 7284 | Dunnett |
|  | *Pik3cg* v3 Saline vs Contra | 0.059 | 3, 16 | 98.53 | <0.0001 | 4081 to 7328 | Dunnett |
|  | *Pten* Saline vs Contra | 1.020 | 3, 16 | 5.512 | 0.0086 | 12794 to 95708 | Dunnett |
|  | *Akt1* Saline vs Ipsi | 0.263 | 3, 16 | 8.548 | 0.0013 | 4693 to 28142 | Dunnett |
|  | *Akt1* Saline vs Contra | 0.136 | 3, 16 | 8.548 | 0.0013 | 7515 to 30964 | Dunnett |
|  | *Akt2* Saline vs Ipsi | 0.035 | 3, 16 | 26.04 | <0.0001 | 6569 to 14824 | Dunnett |
|  | *Akt2* Saline vs Contra | 0.024 | 3, 16 | 26.04 | <0.0001 | 6698 to 14953 | Dunnett |
|  | *Akt3* Saline vs Ipsi | 0.139 | 3, 16 | 32.40 | <0.0001 | 61231 to 165924 | Dunnett |
|  | *Akt3* Saline vs Contra | 0.042 | 3, 16 | 32.40 | <0.0001 | 74049 to 178741 | Dunnett |
|  | *nNos1* Saline vs Ipsi | 0.279 | 3, 16 | 16.04 | <0.0001 | 8185 to 26645 | Dunnett |
|  | *nNos1* Saline vs Contra | 0.144 | 3, 16 | 16.04 | <0.0001 | 11459 to 29919 | Dunnett |
|  | *Jnk3* Saline vs Ipsi | 0.314 | 3, 16 | 34.01 | <0.0001 | 83822 to 192078 | Dunnett |
|  | *Jnk3* Saline vs Contra | 0.172 | 3, 16 | 34.01 | <0.0001 | 112513 to 220769 | Dunnett |
| 2C | *Pik3cg* v3 Saline vs Morphine | 0.519 | 3, 16 | 11.97 | 0.0002 | 34.05 to 5331 | Dunnett |
|  | *Pik3cg* v3 Saline vs Ipsi | 0.067 | 3, 16 | 11.97 | 0.0002 | 2558 to 7855 | Dunnett |
|  | *Pik3cg* v3 Saline vs Contra | 0.057 | 3, 16 | 11.97 | 0.0002 | 2609 to 7906 | Dunnett |
|  | *Akt1* Saline vs Ipsi | 4.011 | 3, 16 | 4.540 | 0.0174 | -4589 to -194.0 | Dunnett |
|  | *Akt3* Saline vs Contra | 0.160 | 3, 16 | 5.733 | 0.0073 | 6062 to 48583 | Dunnett |
|  | *nNos1* Saline vs Ipsi | 0.298 | 3, 16 | 7.532 | 0.0023 | 42.40 to 9644 | Dunnett |
|  | *nNos1* Saline vs Contra | 0.191 | 3, 16 | 7.532 | 0.0023 | 783.0 to 10384 | Dunnett |
|  | *Jnk3* Saline vs Ipsi | 0.356 | 3, 16 | 24.62 | <0.0001 | 12370 to 27812 | Dunnett |
|  | *Jnk3* Saline vs Contra | 0.389 | 3, 16 | 24.62 | <0.0001 | 11355 to 26798 | Dunnett |
| 3A | SC Saline vs Contra | 0.423 | 3, 9 | 14.90 | 0.0008 | 193.3 to 709.2 | Dunnett |
|  | DRG Saline vs Ipsi | 0.334 | 3, 9 | 5.658 | 0.0186 | 32.77 to 203.6 | Dunnett |
|  | DRG Saline vs Contra | 0.466 | 3, 9 | 5.658 | 0.0186 | 9.294 to 180.2 | Dunnett |
| 3B | SC Saline vs Contra | 0.161 | 3, 9 | 10.38 | 0.0028 | 16.04 to 93.28 | Dunnett |
|  | SN Saline vs Ipsi | 8.362 | 3, 9 | 23.73 | 0.0001 | -113.7 to -50.32 | Dunnett |
| 3C | SC Saline vs Ipsi | 7.256 | 3, 12 | 27.22 | <0.0001 | -4.999 to -1.754 | Dunnett |
|  | SC Saline vs Contra | 9.318 | 3, 12 | 27.22 | <0.0001 | -6.113 to -2.868 | Dunnett |
|  | SN Saline vs Morphine | 0.704 | 3, 14 | 22.26 | <0.0001 | 0.02031 to 0.4465 | Dunnett |
|  | SN Saline vs Ipsi | 0.247 | 3, 14 | 22.26 | <0.0001 | 0.3688 to 0.8208 | Dunnett |
|  | SN Saline vs Contra | 0.282 | 3, 14 | 22.26 | <0.0001 | 0.3410 to 0.7931 | Dunnett |
| 4A | Day 1 Vehicle vs Quercetin | 1.205 | 9, 72 | 8.682 | <0.0001 | -3.874 to -0.7635 | Dunnett |
|  | Day 3 Vehicle vs Quercetin | 1.312 | 9, 72 | 8.682 | <0.0001 | -4.816 to -1.706 | Dunnett |
|  | Day 3 Vehicle vs SP600125 | 1.173 | 9, 72 | 8.682 | <0.0001 | -3.358 to -0.2469 | Dunnett |
|  | Day 5 Vehicle vs Thalidomide | 1.682 | 9, 72 | 8.682 | <0.0001 | -5.683 to -2.572 | Dunnett |
|  | Day 5 Vehicle vs Quercetin | 1.990 | 9, 72 | 8.682 | <0.0001 | -7.552 to -4.441 | Dunnett |
|  | Day 5 Vehicle vs SP600125 | 1.984 | 9, 72 | 8.682 | <0.0001 | -7.515 to -4.405 | Dunnett |
| 4B | Day 5 Vehicle vs Thalidomide | 1.558 | 9, 72 | 7.151 | <0.0001 | -6.686 to -2.745 | Dunnett |
|  | Day 5 Vehicle vs Quercetin | 1.834 | 9, 72 | 7.151 | <0.0001 | -9.028 to -5.086 | Dunnett |
|  | Day 5 Vehicle vs SP600125 | 1.646 | 9, 72 | 7.151 | <0.0001 | -7.435 to -3.494 | Dunnett |
| 4D | Day 1 Vehicle vs SP600125 | 1.171 | 9, 72 | 10.16 | <0.0001 | -1.700 to -0.2914 | Dunnett |
|  | Day 3 Vehicle vs SP600125 | 1.356 | 9, 72 | 10.16 | <0.0001 | -2.485 to -1.077 | Dunnett |
|  | Day 5 Vehicle vs Quercetin | 1.492 | 9, 72 | 10.16 | <0.0001 | -2.664 to -1.256 | Dunnett |
|  | Day 5 Vehicle vs SP600125 | 1.740 | 9,72 | 10.16 | <0.0001 | -3.650 to -2.241 | Dunnett |

## Results

### *Decreased mRNA expression of* PI3K, AKT, nNOS, and JNK3 *in the spinal cord, dorsal root ganglia, and sciatic nerves of Morphine tolerant mice after SNL*

Morphine tolerance was confirmed through TPW testing which found significant antinociception on day 1 of behavioral testing in morphine treated mice. But by days 3 and 5, the withdrawal latencies of both vehicle- and morphine-treated mice were not significantly different from each other indicating the development of tolerance by at least day 3 (Fig. 1). MT+SNL mice had lower paw withdrawal thresholds on the ipsilateral versus the contralateral side, as found similarly to previous studies (Kitagawa et al., 2005).

**Figure 1.**
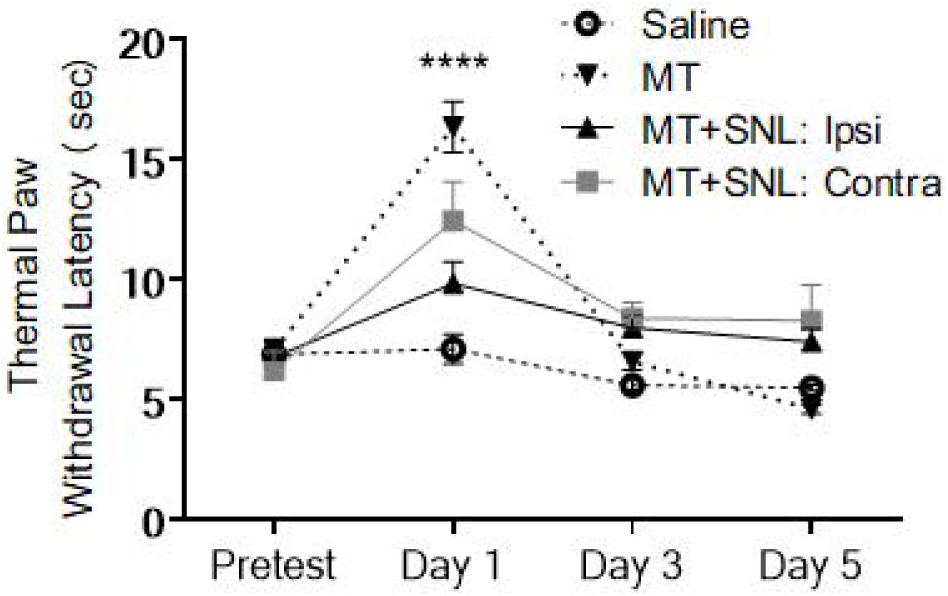
Development of morphine tolerance in mice with or without sciatic nerve ligation. Thermal paw withdrawal thresholds (seconds) in morphine tolerant (MT,15mg/kg, x2 day, 5 days) and vehicle treated mice, and morphine tolerant mice after sciatic nerve ligation (MT + SNL). Withdrawal thresholds measured on both the, ipsi (upward triangles) and contra (squares) sides of MT+SNL mice. Thresholds were measured 30 minutes post injection. Data expressed as mean ± SEM. Two-way ANOVAs with Dunnett post-hoc tests were performed comparing MT mice to saline mice (n=14) and ipsi and contra sides of MT+SNL mice (n=6) for days 1 (including baseline) 3, and 5; ****p<0.0001.

mRNA expression was assessed for 9 different genes in the spinal cord (SC), dorsal root ganglia (DRG), and sciatic nerve (SN) (Fig. 2). Each gene (including different isoforms and splice variants) were chosen because they were all potential downstream targets of the MOP in the PI3K/AKT intracellular signaling pathway (Madishetti et al., 2014). *Pten* is a negative regulator of *Pi3k* (dephosphorylates PIP_3_ to PIP_2_) and *Jnk3* is thought to be upregulated in the spin cord after nerve injury and involved in neuropathic pain (Manassero et al., 2012; Qi et al., 2007) in addition to morphine tolerance in the brain (Fan et al., 2003). Not all of the intracellular mediators within this pathway have been studied with regards to opioid induced tolerance, although previous studies have suggested expression of many of these genes are tissue dependent in the nervous system (Cunha et al., 2010; Liu et al., 2006; Madishetti et al., 2014). The purpose of analyzing gene expression of the MT+SNL mice was to analyze if the addition of a neuropathy model alters gene expression further in mice with morphine tolerance.

**Figure 2.**
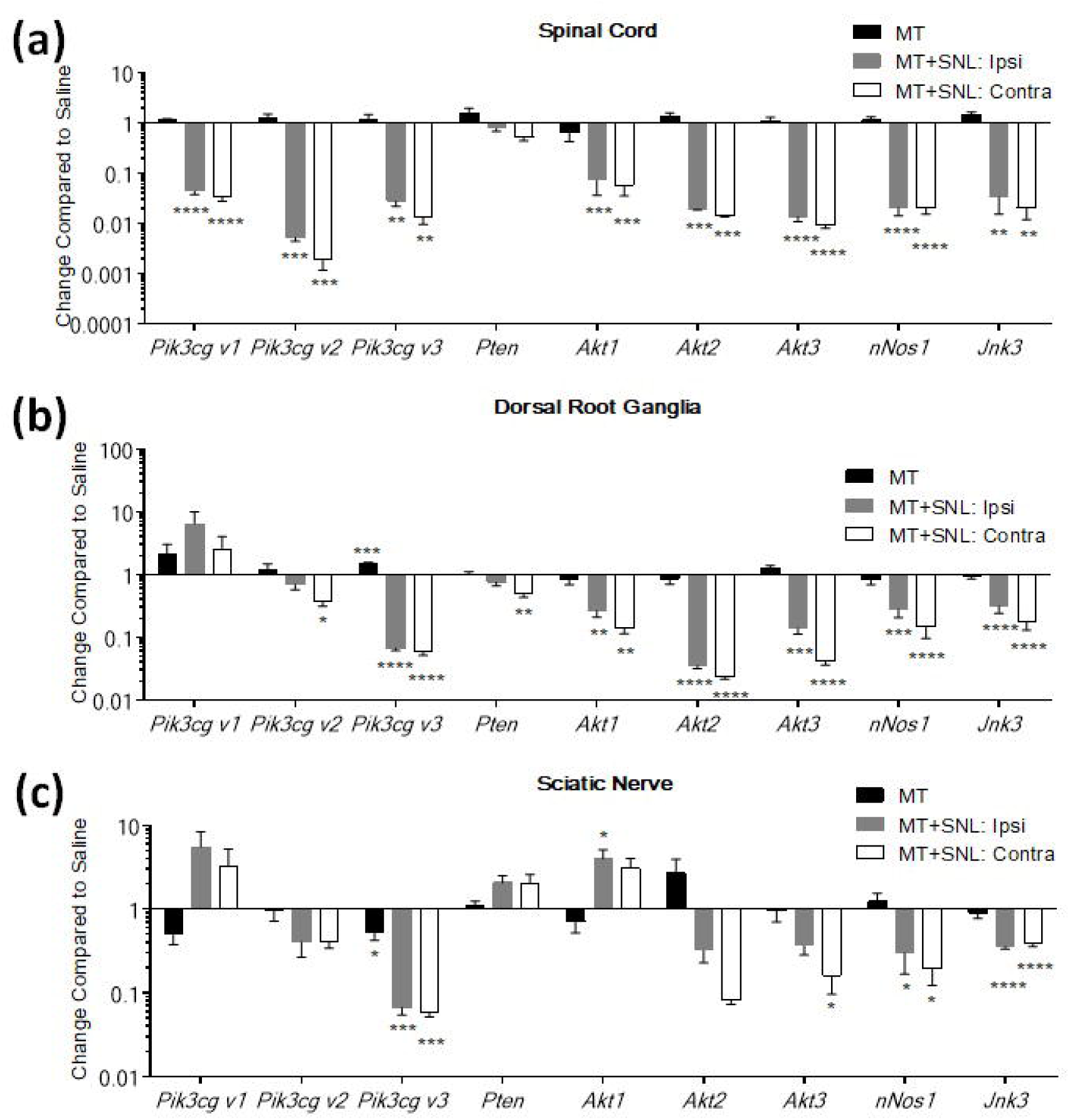
Decreased mRNA expression of the PI3Kγ-AKT-cGMP-JNK pathway in the spinal cord, do rsal root ganglia, and sciatic nerves of morphine tolerant mice after spinal nerve ligation. Fold change of mRNA expression in morphine-treated (MT, 15mg/kg, x2 day, 5 days), and morphine treated mice after SNL (MT+SNL) in the spinal cord (a), DRG (b), and the sciatic nerve (c). The tissues from MT+SNL mice from the injured (i.e. ipsilateral, “ipsi”) and uninjured (i.e. contralateral, “contra”) sides of mice are separated. qPCR analysis data are expressed as fold change over saline-treated mice as mean ± SEM. ANOVA was performed with a Dunnett’s multiple comparisons test comparing mice subcutaneously injected with saline to MT and MT+SNL mice (n=5/group); *p<0.05. **p<0.01; ***p<0.001; ****p<0.0001.

Gene expression within the PI3Kγ-AKT-nNOS-cGMP intracellular signaling pathway was expected to decrease in MT and MT+SNL treatment groups compared to vehicle, while *Pten* and *Jnk3* expression were expected to increase. However, no significant difference was seen in any of the genes analyzed in the spinal cord between MT and vehicle treatment groups (Fig. 2a). And, the only significant change in gene expression seen between MT and vehicle treatment groups was an increase in *Pik3cg v3* in DRG (Fig. 2b) and a decrease in the sciatic nerve (Fig. 2c). For the MT+ SNL treatment group compared to vehicle, there was a significant decrease in gene expression in the spinal cord for all genes except *Pten* (Fig. 2a) with a significant decrease also seen DRG for *Pik3cg v3, Akt1, Akt2, Akt3, Jnk3, and nNos1* (Fig. 2b). Significant decreases in gene expression were also witnessed in the sciatic nerve of MT+SNL mice for *Pik3cg v3, Jnk3, and nNos1* (Fig. 2c). Table 3 provides the baseline expression for each gene and tissue for saline treated mice and a summary of fold change levels is listed in Table 4.

**Table 3.** Average expression of *PI3k* variants, *Akt isoforms*, *nNos1*, and *Jnk3* the spinal cord (SC), dorsal root ganglia (DRG), sciatic nerve (SN) of saline treated mice. Gene expression in # copies. SEM: standard error of the mean.

| Gene | SC Gene Expression | SC ± SEM | DRG Gene Expression | DRG ± SEM | SN Gene Expression | SN ± SEM |
| --- | --- | --- | --- | --- | --- | --- |
| *Pik3cg v1* | 7.55E+02 | 2.96E+01 | 3.60E+01 | 1.33E+01 | 3.84E+01 | 1.37E+01 |
| *Pik3cg v2* | 2.85E+04 | 3.74E+03 | 2.80E+04 | 3.68E+03 | 2.61E+04 | 4.99E+03 |
| *Pik3cg v3* | 3.52E+03 | 2.27E+02 | 6.06E+03 | 8.39E+02 | 5.58E+03 | 1.35E+03 |
| *Pten* | 1.07E+05 | 1.09E+04 | 1.10E+05 | 1.56E+04 | 1.77E+04 | 4.69E+03 |
| *Akt1* | 2.88E+04 | 3.13E+03 | 2.23E+04 | 5.58E+03 | 7.94E+02 | 2.14E+02 |
| *Akt2* | 4.39E+03 | 5.47E+02 | 1.11E+04 | 1.64E+03 | 3.06E+03 | 7.20E+02 |
| *Akt3* | 3.89E+04 | 3.27E+03 | 1.32E+05 | 2.38E+04 | 3.25E+04 | 6.71E+03 |
| *nNOS1* | 2.15E+04 | 2.23E+03 | 2.42E+04 | 2.84E+03 | 6.90E+03 | 1.33E+03 |
| *Jnk3* | 4.37E+04 | 1.16E+04 | 2.01E+05 | 1.35E+04 | 3.12E+04 | 1.60E+03 |

**Table 4.**
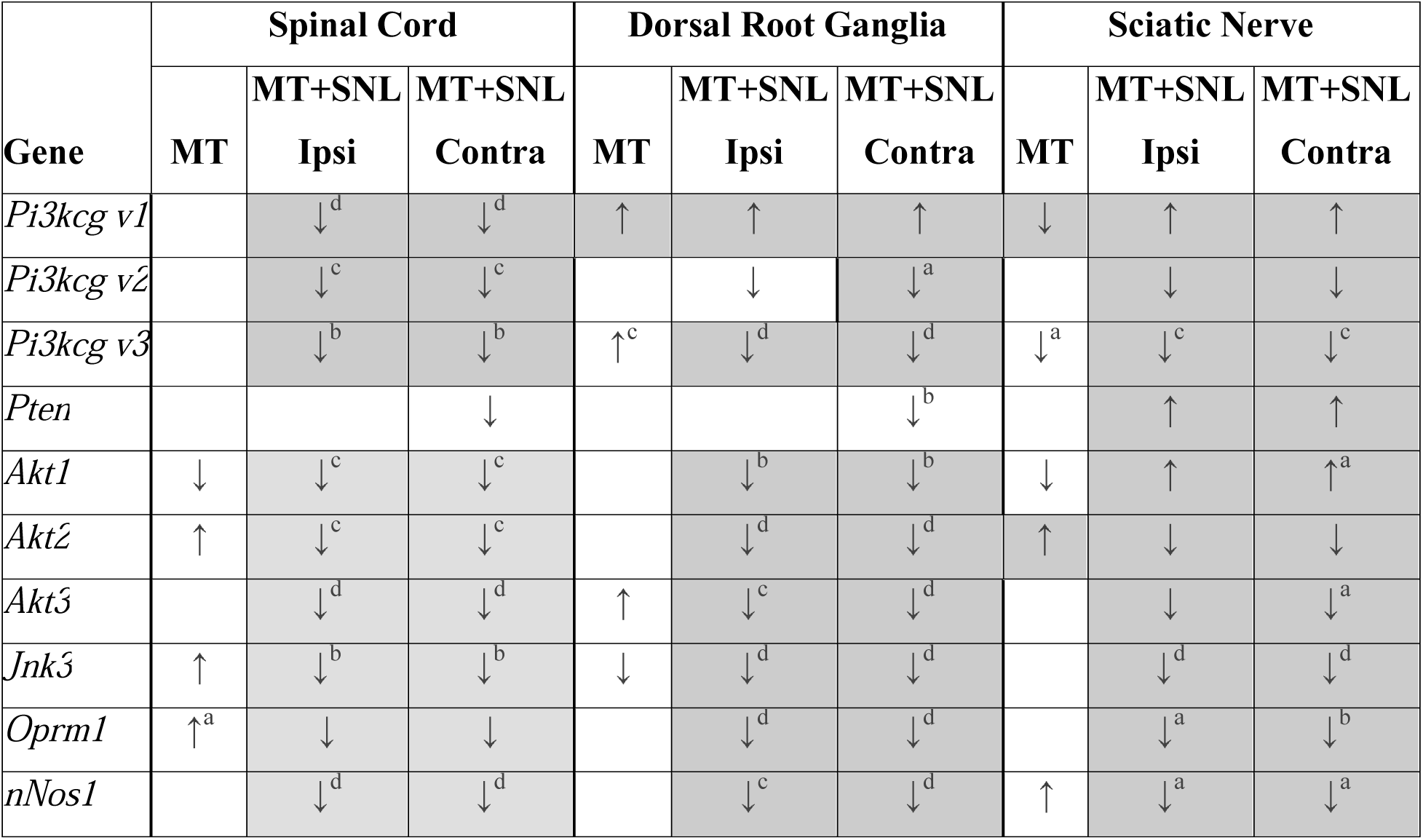
Overview of gene expression changes in MT and MT+SNL treated mice for all tissues. Tissues collected from morphine tolerant mice, and mice after spinal nerve ligation (SNL) including ipsilateral (ipsi) and contralateral (contra) tissues. The arrows indicate an increase (up) or a decrease (down) in gene expression compared to saline treated mice from spinal cord, dorsal root ganglia, and sciatic nerves. Shaded cells indicate at least a two-fold decrease or at least a two-fold increase compared to saline treated mice. ANOVA with Dunnett post-hoc test, ^a^p<0.05; ^b^p<0.01; ^c^p<0.001; ^d^p<0.0001.

| Gene | Spinal Cord |  |  | Dorsal Root Ganglia |  |  | Sciatic Nerve |  |  |
| --- | --- | --- | --- | --- | --- | --- | --- | --- | --- |
|  | MT | MT+SNL<br>Ipsi | MT+SNL<br>Contra | MT | MT+SNL<br>Ipsi | MT+SNL<br>Contra | MT | MT+SNL<br>Ipsi | MT+SNL<br>Contra |
| *Pi3kcg v1* |  | ↓<sup>d</sup> | ↓<sup>d</sup> | ↑ | ↑ | ↑ | ↓ | ↑ | ↑ |
| *Pi3kcg v2* |  | ↓<sup>c</sup> | ↓<sup>c</sup> |  | ↓ | ↓<sup>a</sup> |  | ↓ | ↓ |
| *Pi3kcg v3* |  | ↓<sup>b</sup> | ↓<sup>b</sup> | ↑<sup>c</sup> | ↓<sup>d</sup> | ↓<sup>d</sup> | ↓<sup>a</sup> | ↓<sup>c</sup> | ↓<sup>c</sup> |
| *Pten* |  |  | ↓ |  |  | ↓<sup>b</sup> |  | ↑ | ↑ |
| *Akt1* | ↓ | ↓<sup>c</sup> | ↓<sup>c</sup> |  | ↓<sup>b</sup> | ↓<sup>b</sup> | ↓ | ↑ | ↑<sup>a</sup> |
| *Akt2* | ↑ | ↓<sup>c</sup> | ↓<sup>c</sup> |  | ↓<sup>d</sup> | ↓<sup>d</sup> | ↑ | ↓ | ↓ |
| *Akt3* |  | ↓<sup>d</sup> | ↓<sup>d</sup> | ↑ | ↓<sup>c</sup> | ↓<sup>d</sup> |  | ↓ | ↓<sup>a</sup> |
| *Jnk3* | ↑ | ↓<sup>b</sup> | ↓<sup>b</sup> | ↓ | ↓<sup>d</sup> | ↓<sup>d</sup> |  | ↓<sup>d</sup> | ↓<sup>d</sup> |
| *Oprm1* | ↑<sup>a</sup> | ↓ | ↓ |  | ↓<sup>d</sup> | ↓<sup>d</sup> |  | ↓<sup>a</sup> | ↓<sup>b</sup> |
| *nNos1* |  | ↓<sup>d</sup> | ↓<sup>d</sup> |  | ↓<sup>c</sup> | ↓<sup>d</sup> | ↑ | ↓<sup>a</sup> | ↓<sup>a</sup> |

### Decrease in total JNK while pJNK levels depend on nervous system location in MT+SNL mice

Total and phosphorylated JNK protein concentrations (mg/mL) were analyzed in spinal cord, DRG, and sciatic nerves from vehicle, MT, and MT+SNL treatment groups. Levels of both total and phosphorylated JNK decreased in all three tissues from both MT and MT+SNL treatment groups when compared to the vehicle treatment group (Fig. 3A, B; Table 5) with a significant decrease seen in the contralateral side of the spina cord and the ipsilateral and contralateral side of the DRG (Fig. 4a). Levels of phosphorylated JNK increased in the sciatic nerve and decreased in the DRG for MT and MT+SNL treatment groups compared to vehicle (Fig. 3B).

**Figure 3.**
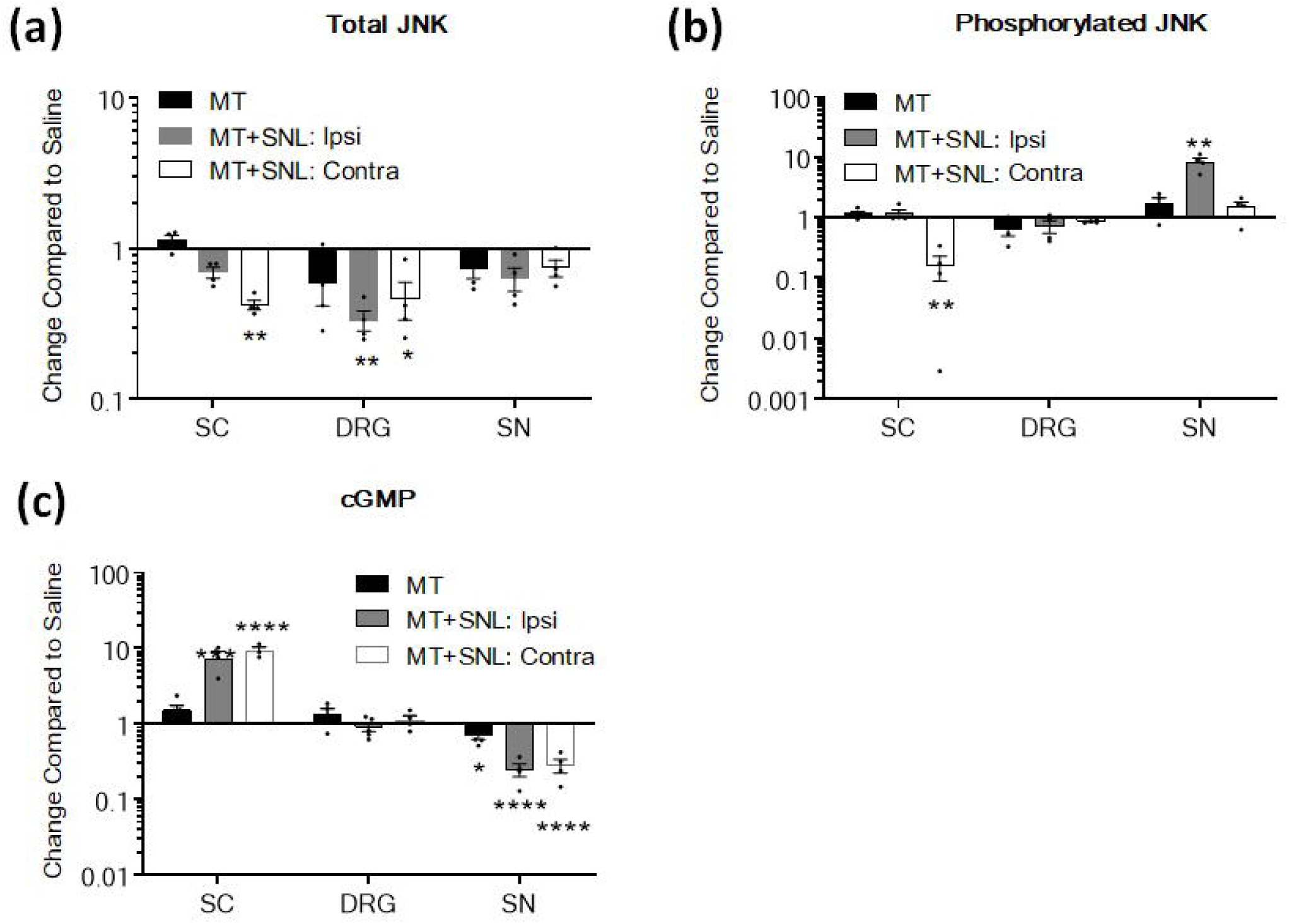
Total and phosphorylated-JNK (pJNK) levels and cGMP levels are altered in the spinal cord (SC), dorsal root ganglia (B, DRG), sciatic nerve (SN) of morphine tolerant mice after spinal nerve ligation. Fold change of morphine-treated (MT, 15mg/kg, x2 day, 5 days), and morphine treated mice after SNL (MT+SNL). The tissues from MT+SNL mice from the injured (i.e. ipsilateral, “ipsi”) and uninjured (i.e. contralateral, “contra”) sides of mice are separated. ELISA analysis of (a)Total JNK (b) pJNK, (c) cGMP levels in MT, saline and MT + SNL SC DRG, and SN tissues. Data are expressed as fold change over saline-treated mice as mean ± SEM. ANOVA was performed with a Dunnett’s multiple comparisons test comparing mice subcutaneously injected with saline to MT and MT+SNL mice (n=5/group); *p<0.05. **p<0.01; ***p<0.001; ****p<0.0001.

**Figure 4.**
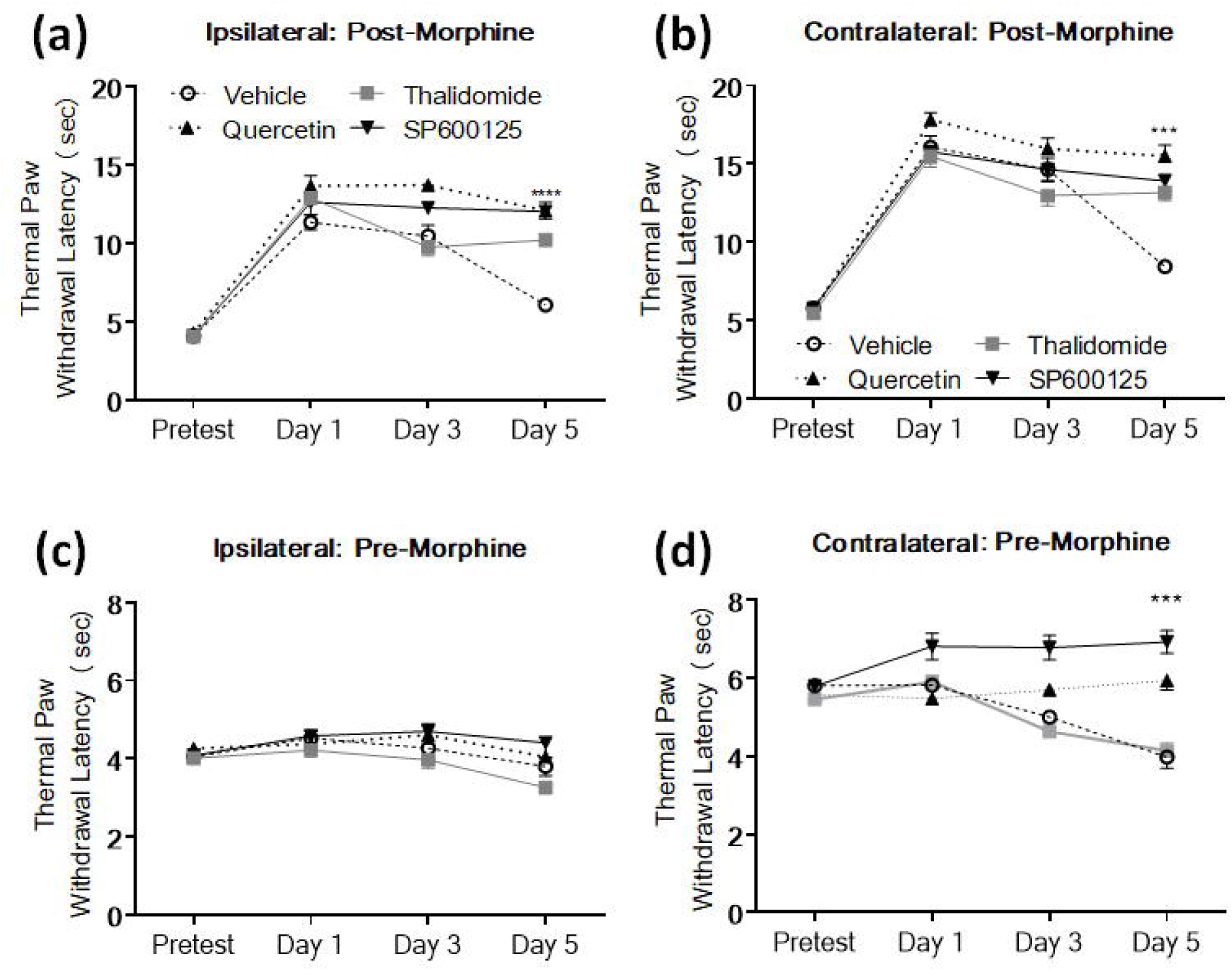
PI3K/AKT pathway inhibitors attenuate the development of morphine tolerance in morphine tolerant mice after spinal nerve ligation. PI3K/AKT pathway inhibitor TPW threshold data for post-morphine treatment of the injured (i.e. ipsilateral, “(a) ipsi and (b) contra sides of MT+SNL mice (15mg/kg, x2 day, 5 day) and pre-morphine, post vehicle or PI3Kcg inhibitor treatment of the (c) ipsi and (d) contra sides of MT+SNL mice. Vehicle was given 30 mins prior to morphine injection), or one of the PI3K/AKT pathway inhibitors: quercetin, thalidomide, or SP600125. Data expressed as mean ± SEM. Repeated measures ANOVA for both ipsi and contra MT+SNL+ inhibitors using a Dunnett post-hoc test to compare back to the vehicle was used for pre- and post-morphine administration (n=5). *p<0.05; **p<0.01; ***p<0.001; ****p<0.0001.

**Table 5.** Total and phosphorylated JNK (pJNK) from ELISA analysis of the spinal cord (SC), dorsal root ganglia (DRG), and sciatic nerve (SN) of saline-treated mice.

| **Tissue** | **JNK Total (mg/mL)** | **JNK Total**<br>**± SEM** | **pJNK (mg/mL)** | **pJNK**<br>**± SEM** |
| --- | --- | --- | --- | --- |
| **SC** | 7.81E+02 | 1.33E+02 | 6.51E+01 | 1.38E+01 |
| **DRG** | 1.77E+02 | 2.02E+01 | 2.43E+01 | 4.93E+00 |
| **SN** | 9.71E+01 | 2.17E+01 | 1.11E+01 | 2.44E+00 |

### cGMP levels increase in the spinal cord and decrease in the sciatic nerve of MT+ SNL mice

cGMP nucleotide concentrations for the MT and MT + SNL mice were analyzed for spinal cord, dorsal root ganglia, and sciatic nerve and compared to saline mice (Fig. 3C; Table 6). cGMP was chosen for analysis as it is responsible for activating protein kinase G (PKG), which phosphorylates and activates K_ATP_ channels either directly or through the ROS/calmodulin/CaMKII signaling cascade (Cao et al., 2016; Chai et al., 2011; Ding et al., 2017). We hypothesized cGMP nucleotide levels would decrease in MT and MT+SNL mice as a decrease in cGMP would lower PKG activation, subsequently lowering K_ATP_ channel phosphorylation and activation. There was no significant increase or decrease in cGMP protein levels for the spinal cord, dorsal root ganglia, and sciatic nerve (p>0.05) of MT mice without nerve injury. Significant increases in cGMP were found in the spinal cord of MT+ SNL mice, while significant decrease was found in the sciatic nerves.

**Table 6.**
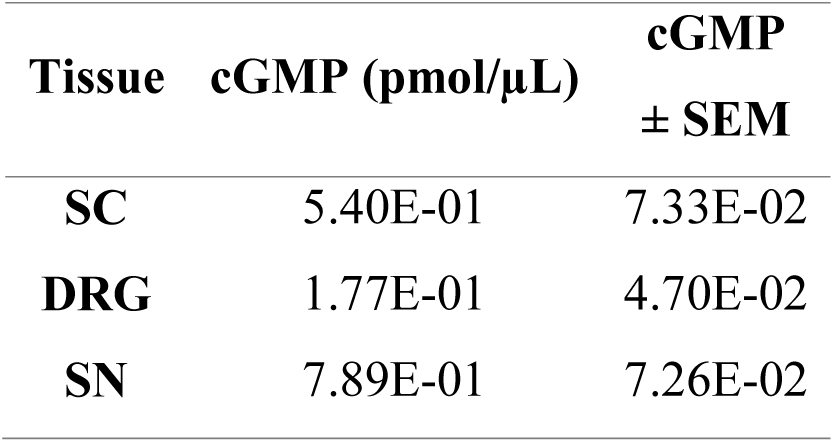
cGMP nucleotide levels from ELISA analysis of the spinal cord (SC), dorsal root ganglia (DRG), and sciatic nerve (SN) of saline-treated mice.

### *Pharmacological inhibition of the* PI3Kγ-AKT-nNOS-cGMP *pathway attenuates morphine antinociception*

To determine if the inhibition of the PI3Kγ-AKT-nNOS-cGMP pathway would alter the development of opioid induced tolerance, we systemically administered either thalidomide, quercetin, or SP600125 before morphine in the MT+SNL treatment group and TPW was tested. We hypothesized inhibiting the PI3Kγ-AKT-cGMP-K_ATP_ pathway would decrease or attenuate opioid induced tolerance for thalidomide or quercetin as they would decrease PI3Kγ -AKT activity. Additionally, we hypothesized SP600125 would attenuate morphine tolerance as the inhibition of the phosphorylation of JNK could potentially drive PI3Kγ-AKT-nNOS-cGMP pathway to phosphorylate and open K_ATP_ channels. Injured (i.e. ipsilateral, “ipsi”) and uninjured (i.e. contralateral, “contra”) sides of the MT+SNL mice were compared after administration of thalidomide, quercetin, SP600125, and the vehicle (Fig. 4). On day five of testing, thalidomide, quercetin, and SP600125 treatment groups had significantly increased paw withdrawal latencies compared to vehicle (Fig. 4a). On the contralateral side, no significant differences were seen on days one and three compared to the vehicle (Fig. 4b). On day five, however, thalidomide, quercetin, and SP600125 had significantly increased paw withdrawal latencies compared to vehicle after morphine administration (Fig. 4b). Mechanical paw withdrawal thresholds were also significantly increased after quercetin and SP600125 administration (before morphine) after five days of testing, but not after thalidomide administration (Fig. 4d).

## Discussion

In this study, differences in gene expression for the PI3Kγ-AKT-cGMP-JNK signaling pathway were found in the spinal cord and in the DRG and sciatic nerves during morphine tolerance with underlying spinal nerve injury (Figure 5). Phosphorylated levels of JNK and cGMP were decreased in the spinal cord but increased in the sciatic nerve. Inhibition of the PI3Kγ-AKT-cGMP-JNK signaling pathway using a pharmacological strategy attenuated the development of morphine tolerance in mice. Furthermore, continued activation of opioid receptors by morphine after SNL decreased *PI3k*, *Akt*, and *Jnk3* gene expression and increased cGMP nucleotide levels in the spinal cord. Continued activation also decreased *PI3k*, *Akt*, and *Jnk3* gene expression and decreased cGMP nucleotide levels in the PNS, but increased p-JNK.

**Figure 5.**
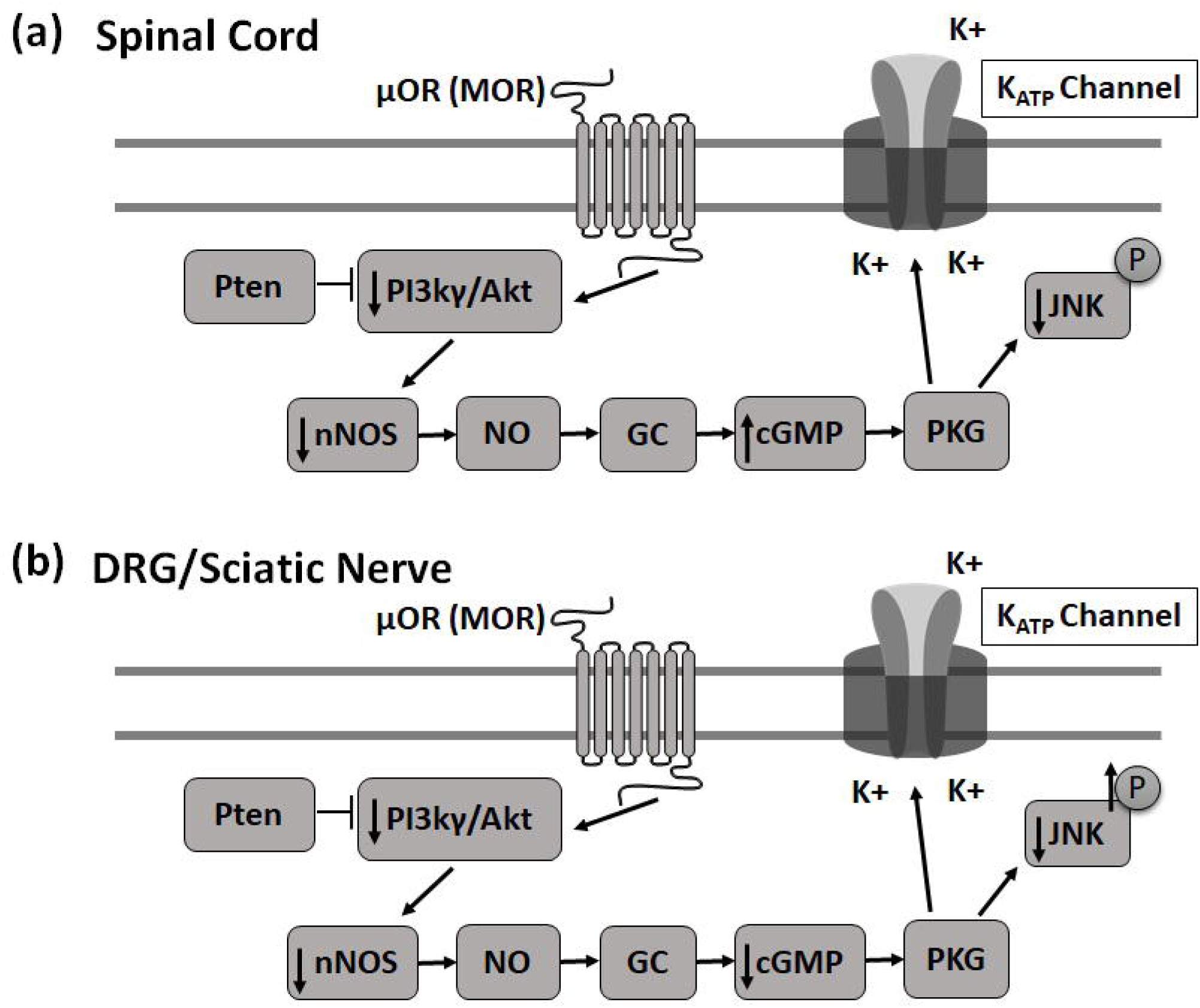
Schematic representation of a possible molecular basis of opioid induced tolerance in morphine tolerant mice after spinal nerve ligation. Continued activation of opioid receptors (a) decreased gene expression of *PI3k*, *Akt*, *nNos*, and *Jnk3* and increased cGMP nucleotide levels in the SC, and (b) decreased gene expression of *PI3k*, *Akt*, *nNOS*, and *Jnk3* and decreased cGMP nucleotide levels, and increased p-JNK in the PNS. The increased phosphorylation of JNK seen in the PNS is facilitated by the PI3Kγ-AKT-cGMP pathway, lowering the activity of K_ATP_ channels or effecting other downstream pathways.

Repeated subcutaneous injections of morphine decreased *Pik3cg*, *Akt1, Akt2,* and *Akt3* gene expression in the spinal cord, DRG, and sciatic nerves of MT+SNL mice. PI3K and its downstream target, Akt, are involved in the modulation of nociceptive information, such as neuropathic pain (Xu et al., 2019) and morphine tolerance (Chen et al., 2018; König et al., 2010). Some of the results presented here were inconsistent with other reports indicating an increase in *Pik3*/*Akt* expression in the sciatic nerve of mice after SNL (Murashov et al., 2001) and in the DRG after oxaliplatin administration (Jiang et al., 2016). One important observation was the differences in *Pik3cg* variant gene expression across different tissue types. Overall, gene expression of *Pik3cg* variants 2 and 3 were higher than *Pik3cg* variant 1 for all tissue types, but *Pik3cg* variant 3 was more abundant in the PNS than in the spinal cord. The *Pik3cg* variant 3 was significantly decreased in all three tissue types compared to saline-treated mice, indicating that different splice variants are affected differently under pain conditions.

Recent studies have found acute morphine treatment can induce MOP dependent stimulation of AKT in DRG (Madishetti et al., 2014), but studies investigating *PI3k* and *Akt* expression during *in vivo* chronic morphine administration are sparse. The PI3K/Akt proteins are involved in the initiation of multiple intracellular pathways during chronic pain, including activation of mammalian target of rapamycin (mTOR) and neutrophilic alkaline phosphatase 1 (NALP1) (Chen et al., 2017). Further downstream mediators are implicated in microglia migration and activation during neuropathic pain and chronic morphine administration (Guo et al., 2017; Horvath and DeLeo, 2009; Xu et al., 2014). Three different isoforms of *Akt* are expressed in the nervous system, and both *Akt2* and *Akt3*are involved in MOP signaling (Cho et al., 2001; Kim et al., 1999). Comparing the baseline expression of the various *Akt* isoforms, *Akt3* was the most abundant in the tissues used in this study. In the ipsilateral sciatic nerve of MT+SNL mice, *Akt1* was >10 fold higher than *Akt2* and *Akt3*. A previous study in *Drosophila*suggests *Akt1* is upregulated around the site of nerve injury and is involved in the removal of axonal debris (Musashe et al., 2016), and the increased expression of *Akt1* found in the MT+SNL mice would match this observation.

*NO* and *nNos* are extensively studied for their roles in nociception (Miclescu and Gordh, 2009), but there is a lack of consensus between their roles in the CNS and PNS under chronic opioid exposure. Data presented here indicates *nNos* decreases in MT+SNL mice, consistent with a previous study in rats finding *nNos* activity decreases in the spinal cord after SNL (Yang et al., 2007). In a mouse model of peripheral nerve injury, the expression of nNOS within DRG was increased, but the expression of nNOS within the spinal cord was unchanged (Guan et al., 2007). These findings suggest nNOS plays a critical role in hypersensitivity during chronic pain, but is location dependent. Increased *nNos* activity is also associated with opioid-induced tolerance (Lutz et al., 2015), as the inhibition of *nNos* can attenuate the development of morphine tolerance in spinal microglia (Liu et al., 2006). Morphine stimulates the PI3K/AKT pathway, activating NO and subsequently PKG, which then triggers K_ATP_ channels to open (Cunha et al., 2010). However, NO can directly activate K_ATP_ channels in the presence of guanylyl cyclase and PKG inhibitors (Kawano et al., 2009b), indicating not all of the components involved in the PI3Kγ-AKT-cGMP pathway are necessary for K_ATP_-induced analgesia. Data presented here suggest the activity of this pathway is tissue-dependent as a marked decline in PI3Kγ-AKT-nNOS gene expression in the spinal cord and DRG of MT+SNL mice was found, but not in the sciatic nerve.

cGMP nucleotide levels were found to be significantly lower for MT mice than saline-treated mice in the sciatic nerve. Conversely, during *in vitro* morphine exposure, cGMP levels have been found to be increased in tuberomammillary nucleus neurons of neonatal rats (Mo et al., 2005). Our data suggest that morphine tolerance, with or without nerve injury, results in a decrease of cGMP nucleotide levels in the peripheral nervous system. Increased cGMP was found in the spinal cord of MT+SNL treated mice, consistent with previous studies finding chronic sciatic nerve constriction injury increased spinal cGMP levels in rats (Siegan et al., 1996) and in mice after chronic morphine exposure (Liang and Clark, 2004). Previous data indicate JNK is upregulated in the spinal cord after SNL in rats (Zhuang et al., 2006) and contributes to inflammatory responses (Nomura et al., 2013). Mice lacking JNK1, JNK2, or JNK3 also have attenuated morphine tolerance compared to wild-type mice (Hervera et al., 2012). Additionally, *Jnk3* is upregulated during morphine tolerance in the frontal cortex of rats (Fan et al., 2003) and p-JNK is upregulated after ultra-low dose morphine administration in the spinal cord (de Freitas et al., 2019), thalamus, prefrontal cortex, and periaqueductal grey of mice (Sanna et al., 2014). Overall, there was no observation of increased *Jnk3* expression in the spinal cord, DRG, or sciatic nerve for any of our treatment groups. Total JNK was downregulated across all tissues, but p-JNK was upregulated only in the sciatic nerve of MT+SNL mice, similar to a previous report indicating increased p-JNK expression in DRGs after morphine exposure over six days (Ma et al., 2001). The differences between cGMP and JNK activity in the CNS and PNS are important, as they once again highlight the involvement of potentially different intracellular mechanisms occurring in the nervous system during chronic pain and opioid signaling.

Quercetin, thalidomide, and SP600125 all attenuated the development of morphine tolerance in SNL mice. Quercetin, a PIK3cg inhibitor, previously has shown to reduce thermal hyperalgesia in diabetic mice (Anjaneyulu and Chopra, 2003). The thalidomide and SP600125 results presented here are supported by previous studies using rodent models of opioid tolerance (Babbini and Davis, 1972; Chai et al., 2011; Khan et al., 2017; Yuill et al., 2016). Thalidomide is also involved with the inhibition of TNF-α, a mediator of the inflammatory immune response (Klausner et al., 1996), which is important to note, as ligation of the spinal nerve creates a local inflammatory response contributing to the development of tolerance and/or mechanical hyperalgesia. Opioid tolerance and hyperalgesia share common cellular mechanisms, as similar manipulations can block both tolerance and hyperalgesia (Chen et al., 2008; King et al., 2005; Lee et al., 2011; Vanderah et al., 2001) and the behavioral findings presented here support these observations.

The activation of K_ATP_ channels to produce analgesic effects are readily reported (Afify et al., 2013; Xia et al., 2014), yet many of the intracellular signaling pathways involved in K_ATP_ channel activation are still largely unknown (Cao et al., 2016). The increased phosphorylation of JNK seen in the PNS could lower the activity of K_ATP_ channels (Hervera et al., 2012). Alternatively, the loss of K_ATP_ channel activity during chronic pain (Luu et al., 2019; Zoga et al., 2010) and/or chronic opioid administration (Wu et al., 2011) could increase JNK activation by decreased suppression of p-JNK. This idea is supported in a model of post-surgical pain in rats, where administration of a K_ATP_ channel agonist attenuated mechanical hypersensitivity (Zhu et al., 2015) and a in model of morphine tolerance in mice, by decreasing p-JNK expression (Cao et al., 2016). The initial decreased K_ATP_ channel expression and function in the spinal cord could be explained by various other signaling molecules, including cAMP and calmodulin (Kawano et al., 2009a).

Overall, results presented here suggest a link between expression of the PI3Kγ-AKT-cGMP-JNK signaling pathway and the development of morphine tolerance in nerve-injured mice. Further investigation of the PI3Kγ-AKT-cGMP-JNK signaling pathway will likely provide novel targets for preventing and/or treating chronic morphine tolerance and other related clinically relevant conditions such as opioid-induced hyperalgesia and withdrawal.

## Original Article

### Funding Sources

Funding provided by the NIH to A.H.K. (K01 DA042902), the University of Minnesota Integrated Biosciences Graduate Program to T.O., and the University of Minnesota Duluth Undergraduate Research Opportunity Program award to T.J.

### Significance

These findings provide novel evidence of the differences in PI3Kγ-AKT-cGMP-JNK signaling pathway involvement during morphine tolerance in the central versus peripheral nervous system. These data also confirm that systemic delivery of agents that reduce JNK activity can attenuate the development of morphine tolerance in mice with underlying neuropathic pain. Together the results here are translationally relevant and provide novel targets for preventing and/or treating chronic morphine tolerance associated with chronic pain treatment employing opioids.

## Acknowledgements

The authors thank Kayla Johnson for reading an earlier version of this manuscript and James Bjork for assistance with primer design.

## Author Contributions

AHK and TO conceived the idea and contributed to the study design. TO, TJ, and MM acquired the data. TO performed the statistical analysis. AHK and TO interpreted the data, drafted the paper, and revised the article critically for intellectual content. All authors discussed the results and commented on the manuscript.

